# Combining weather patterns and cycles of population susceptibility to forecast dengue fever epidemic years in Brazil: a dynamic, ensemble learning approach

**DOI:** 10.1101/666628

**Authors:** Sarah F. McGough, Cesar L. Clemente, J. Nathan Kutz, Mauricio Santillana

## Abstract

Transmission of dengue fever depends on a complex interplay of human, climate, and mosquito dynamics, which often change in time and space. It is well known that disease dynamics are highly influenced by a population’s susceptibility to infection and microclimates, small-area climatic conditions which create environments favorable for the breeding and survival of the mosquito vector. Here, we present a novel machine learning dengue forecasting approach, which, dynamically in time and adaptively in space, identifies local patterns in weather and population susceptibility to make epidemic predictions at the city-level in Brazil, months ahead of the occurrence of disease outbreaks. Weather-based predictions are improved when information on population susceptibility is incorporated, indicating that immunity is an important predictor neglected by most dengue forecast models. Given the generalizability of our methodology, it may prove valuable for public-health decision making aimed at mitigating the effects of seasonal dengue outbreaks in locations globally.

## Introduction

Due to emerging sensor technologies and computational advances, the last decade has seen significant strides in the way data is generated and collected, resulting in large volumes of complex information known as “Big Data.” The recent availability of these data has opened up the possibility of new and complementary avenues for epidemic monitoring that leverage satellite imagery^1,2^, Internet search engine activity^3,4^, social media^5^, mobile phones^6,7^, genomics^8,9^, and disease surveillance databases^10,11^. This has opened up opportunities to posit and explore more hypotheses for characterizing the causes and outcomes of disease transmission, population behavior, environmental conditions, and other potential indicators of population health. Exploiting these relationships to generate reliable prospective forecasts would benefit health systems by allowing early mobilization of resources for the prevention of morbidities and deaths in the face of public health threats. A major challenge in disease forecasting is developing algorithms that can autonomously and continuously learn from these complex and ever changing dynamical systems, uncovering patterns and signals with little human effort. Machine learning algorithms are ideally suited for such unsupervised or semi-supervised tasks. Indeed, they are having profound impact across a wide range of application fields due their ability to aid in learning and discovery.

One such complex system is the interplay of human, climate, and mosquito dynamics that give rise to the transmission of mosquito-borne diseases like Dengue. Dengue fever, a viral mosquito-borne disease transmitted predominately by the *Aedes aegypti* and *Aedes albopictus* mosquitoes, infects an estimated 390 million people per year, with nearly half the world’s population living at risk of infection^12^. The global burden of dengue has doubled every 10 years over the last 3 decades^13^, and the disease is projected to expand its latitude range as global temperatures increase and create new suitable habitats for the *Aedes* mosquitoes among previously-unexposed human populations^14^. Short-term climate conditions, particularly temperature and precipitation, can create favorable conditions for the breeding and survival of *Aedes* mosquitoes that may increase the transmission of the dengue fever virus in humans. Distinct ranges of temperature and precipitation have been observed to have an influence on the extrinsic incubation period^15,16^, mosquito maturation rate^17^, length of larval hatch time^18^, survival rate^19^, and biting rate^20^. However, the relationships that govern these parameters and give rise to dengue transmission are complex and dynamic, changing over time and across geographies. Moreover, multi-year cycles of dengue fever outbreaks, caused by one or more circulating dengue fever serotypes (DENV I, II, III, IV) and short-term immunity conferred after infection, add an important layer of complexity to prediction^21^.

The dengue forecasting literature lacks a systematic, self-adaptive, and generalizable framework capable of identifying weather and population susceptibility patterns that may be predictive of dengue fever outbreaks, particularly at the city-level. Vector-borne diseases commonly exhibit spatial heterogeneity, a result of spatial variation in vector habitat, weather patterns, and human control actions^22–25^. For developing forecast systems, this feature implies a trade-off between model consistency and spatial resolution. As a consequence, most studies to date focus on producing ad-hoc predictions for a single location, ranging from the national-to the city-level^26–28^, while others build and evaluate multiple modeling strategies per study site in efforts to manually identify relationships between weather patterns and dengue incidence over different geographies and temporal windows^29,30^. Both approaches highlight the difficulty in producing forecast models that are viable in diverse settings. In contrast, data-driven techniques demonstrate promise by learning from multi-scale, complex systems and automatically adapting to new information, i.e. they provide an unsupervised or semi-supervised machine learning infrastructure. A recent descriptive study showed the promise of a data-driven approach in identifying weather patterns with meaningful signals for dengue fever outbreaks^31^. Specifically, their data-driven strategy identified temperature and frequency of precipitation as key features in forecasting dengue outbreaks by extracting windowed time intervals for different cities that were highly predictive. Motivated by such unsupervised learning algorithms, we build upon this data-driven strategy to build a richer forecasting algorithm.

### Our contribution

Focusing on the important but complex relationship between dengue incidence and (a) weather patterns and (b) the empirically observed 3-4 year disease burden cycles, we present a data-driven, machine learning approach capable of autonomously and continuously identifying weather patterns and cycles in population susceptibility to predict dengue fever outbreak years. Specifically, our approach is *dynamic*, since it automatically identifies emerging predictive patterns as new information becomes available; it is adaptable to *multiple and heterogeneous* study areas with *high spatial resolution*, as it leverages a *publicly and globally available* meteorological dataset. We show the predictive potential of this framework in 20 Brazilian municipalities by producing annual retrospective, out-of-sample epidemic forecasts at the city-level, months ahead of the historically-observed seasonal onset of dengue epidemics. We assess the feasibility of this autonomous learning approach using two simple weather inputs (temperature, rainfall) in 20 Brazilian cities with diverse microclimates and diverse Dengue incidence seasonal patterns, and, to give transparency to our framework, attempt to characterize the conditions under which predictions are unsuccessful.

## Results

### Exploiting weather signals to create a data-driven forecast system

We obtained data on both annual dengue fever cases (Brazilian Ministry of Health) for 2001-2017 and on daily temperature and precipitation (GMAO-NASA) for 2000-2016, for 20 dengue-endemic municipalities (Fig. 1, Table S1) in Brazil. Weather patterns were extracted and analyzed across hundreds of partially-overlapping time intervals collectively spanning the last 7 months of a given year, a time period that typically precedes the onset of epidemic outbreaks in Brazil. Each of these patterns were then assessed for their ability to predict an outbreak year (defined as a year in which the number of cases exceeds 100 per 100,000 persons) for the subsequent year. Retrospective and fully out-of-sample forecasts, trained on a yearly-expanding window, were produced for 10 years (2008-2017) and for each time interval using support vector machines, a binary classifier. Every year, the time intervals with high historical predictive power were automatically selected and evaluated in the upcoming year to produce out-of-sample predictions for the subsequent dengue season (Fig. 1). An ensemble approach was then implemented to determine, in an completely out-of-sample fashion (using the first 4 years of out-of-sample predictions to inform ensemble model selection), whether a year would be epidemic or not for the next 6 years (2012-2017).

**Figure 1.**
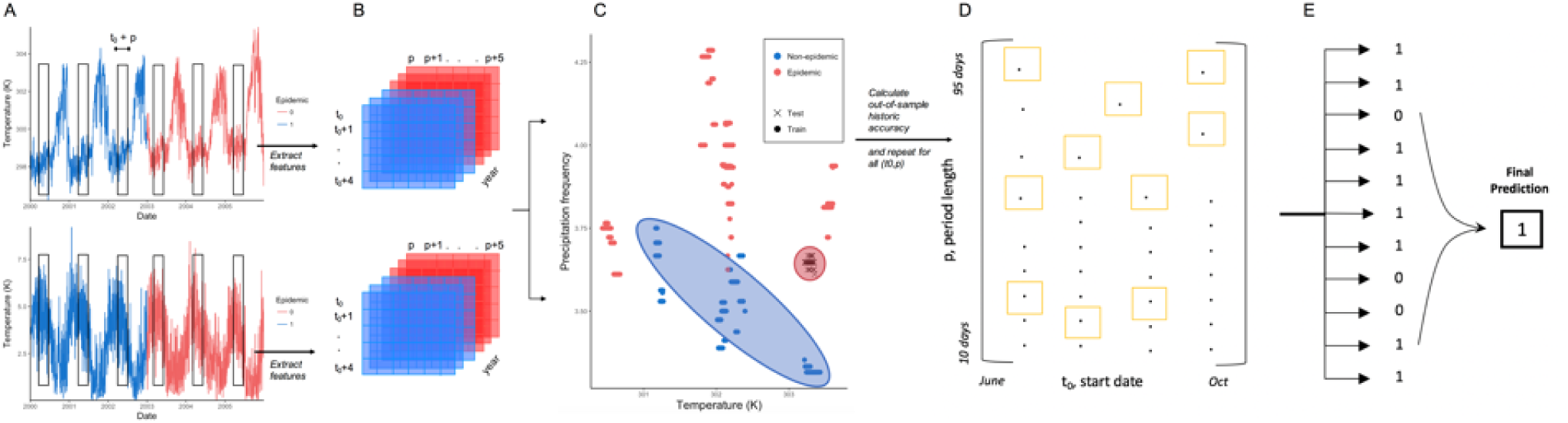
Ensemble forecast workflow. (A) To predict next year’s epidemic status, we extract features from a daily time series of temperature (K) and precipitation (mm) over a defined (t_0_, p) time interval and for each year in the training period. (B) We produce an array of features corresponding to the mean value of temperature and precipitation over the (t_0_, p) interval, and (C) train a support vector machine to classify next year’s epidemic status. (D) This process is repeated for all 432 (t_0_, p) intervals, and the top 11 models are automatically selected to (E) contribute to a majority voting system based on historic out-of-sample accuracy.

This system, which autonomously identifies and exploits the predictions of multiple time windows during the calendar year, makes it possible to identify temporally similar regions of highly predictive periods of the year preceding dengue outbreaks, here referred to as “weather signatures.” Weather signatures represent time windows across years that show strong influence (predictive power) on the incidence of dengue in a subsequent year. We observed that cities where our methodology lead to higher prediction accuracy tended to have clear and robust weather signatures over the years, while cities where our approach was not strongly predictive did not exhibit consistent and robust weather patterns (Figs. 2, 3A). Further, we observed that strong weather signatures in our sample of cities often corresponded with or preceded important alternating tropical seasons, such as rainy and dry seasons.

**Figure 2.**
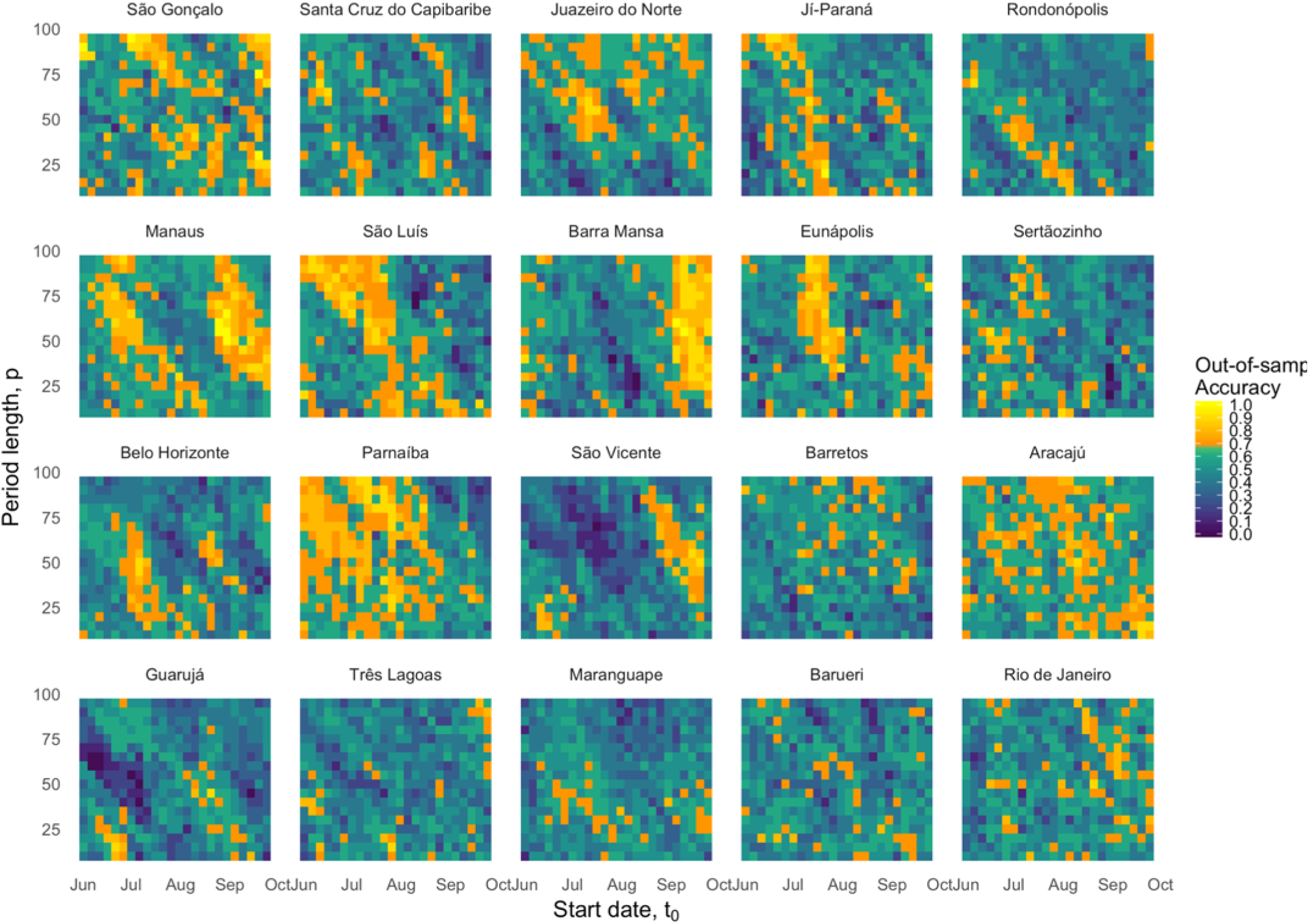
The 10-year (2008-2017) out-of-sample forecast accuracy (%) for each time window of temperature and precipitation, by municipality. The x-axis (t_0_) indicates the start date of the time interval, and the y-axis (p) indicates the length of the time interval from which weather data were gathered (10-95 days). Models achieving at least 7/10 correct out-of-sample forecasts are shown in shades of yellow. Municipalities are ordered by decreasing ensemble prediction accuracy; that is, the proportion of years correctly forecasted by the ensemble method over the years 2012-2017.

**Figure 3.**
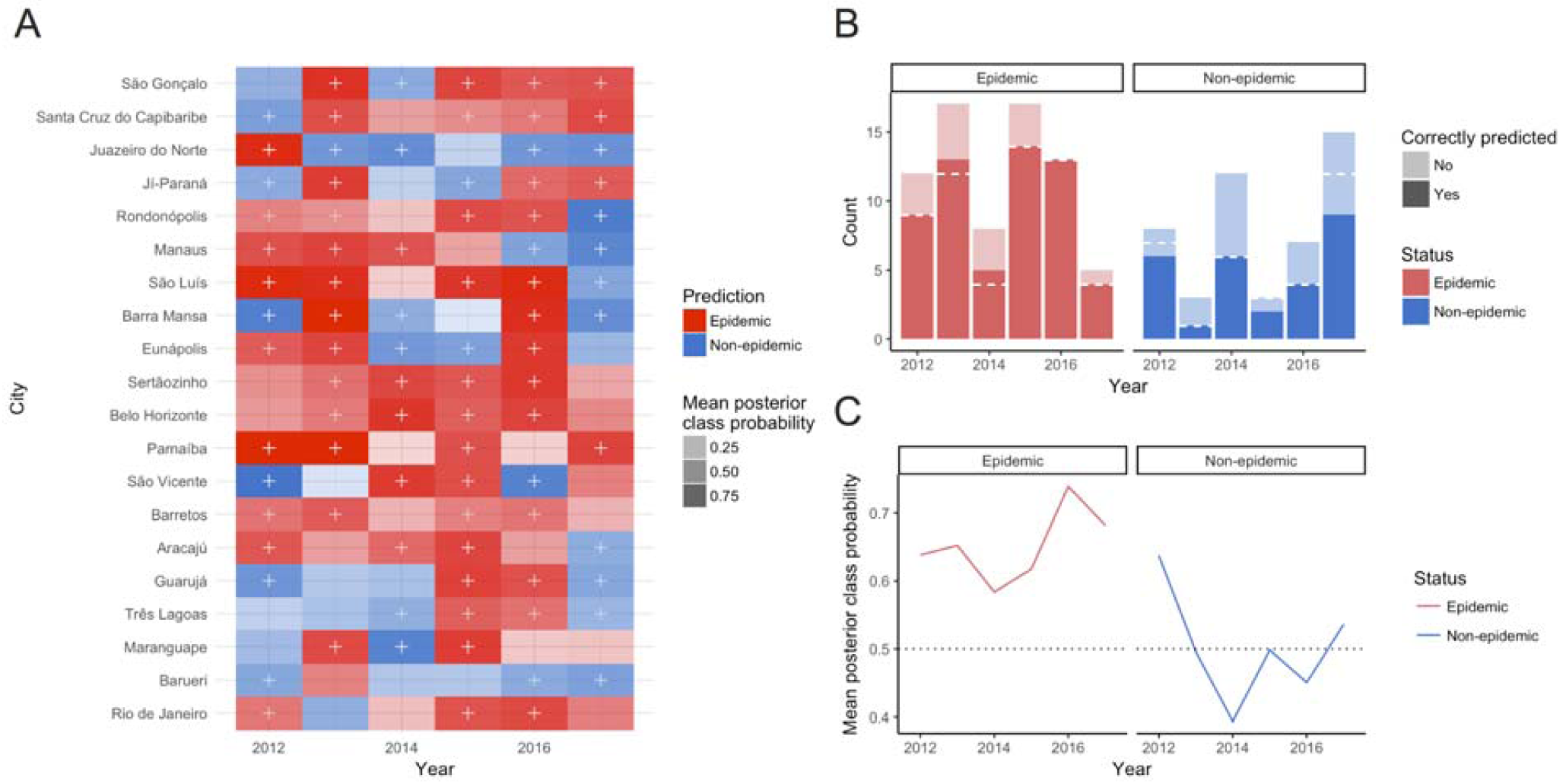
Weather-based prediction results for 120 municipality-years. A) Annual out-of-sample forecasts of outbreak status (epidemic/non-epidemic) for 20 Brazilian municipalities from 2012-2017, shaded by mean posterior probability (MPP) of the true outbreak status. Correct forecasts are indicated by a plus (+) sign, and cells with light shading indicate that the model predicted the correct class with low probability. Municipalities are ordered by decreasing ensemble prediction accuracy; that is, the proportion of years correctly forecasted by the ensemble method over the years 2012-2017. B) The number of total epidemic and non-epidemic years correctly forecasted across 20 municipalities, by year. The dashed white line indicates the number correctly forecasted after incorporation of empirically-observed dengue cycles. C) The mean posterior class probability across municipalities, by year and epidemic status.

### Weather-based forecasting performance

Using weather data (temperature and frequency of precipitation) alone to predict annual dengue outbreaks, our approach correctly forecasted 81% of all epidemic years across 20 municipalities in Brazil between 2012-2017 (Table 1, Fig. 3). For reference and as a baseline, the frequency of epidemic and non-epidemic years was 60% and 40%, thus, a naive approach that predicts that all years are epidemic (the class majority) would achieve an overall accuracy of 60%. Our approach only identified 58% of non-epidemic years correctly. This resulted in an *overall accuracy* of approximately 72%. Our approach significantly exceeded (p=0.005) the predictive power of a naive predictor.

**Table 1.**
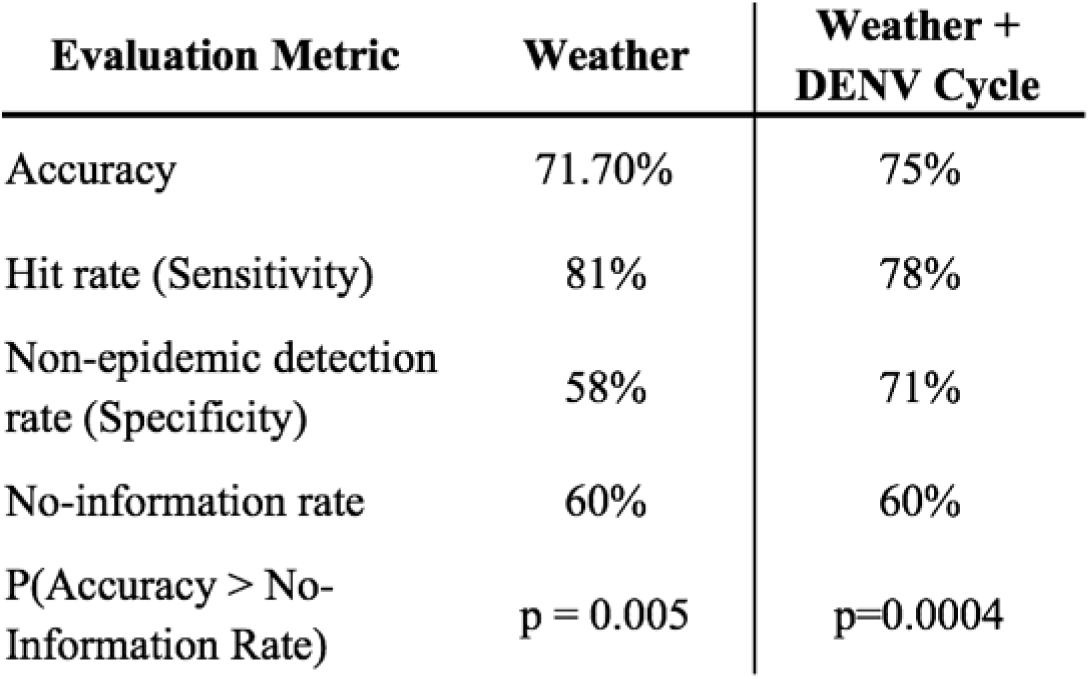
Performance of weather-based out-of-sample forecasts across 120 municipality-years in Brazil, with and without consideration for DENV susceptibility cycles.

### Incorporating empirically observed dengue susceptibility cycles

The previously described weather-based ensemble approach ignores important factors that may influence the emergence of epidemic outbreaks from year to year, such as the population susceptibility to being infected with the virus. Specifically, endemic transmission of dengue fever is typically distinguished by periodic outbreak cycles of around 3-4 years. These outbreak cycles are thought to occur as a result of 1) an exhaustion of the susceptible population after an outbreak, and 2) and short-term cross-immunity to other circulating DENV serotypes after infection^21^, although can also be complicated by increased severity of a second infection^32^. Both factors result in a depletion of the population vulnerable to infection, and act as barriers to subsequent outbreaks. Independent of climate variability over the years, we expect some preservation of these susceptibility cycles.

Inspired by this phenomenon, we implemented a data-driven hidden Markov model by empirically computing the frequency of transitioning between multiple sequences of epidemic and non-epidemic years. If the weather-based approach led to a very unlikely prediction, given the previously-observed sequence of outbreaks (for example, predicting an outbreak year when the susceptible population may be depleted as a consequence of multiple prior years of high Dengue activity), a decision rule was implemented to automatically override the weather-based prediction with a more likely prediction produced using the hidden Markov model (described in detail in *Supplementary Material)*.

### Combining dengue cycles with weather patterns improve forecasts

Compared to the exclusively weather-based approach, incorporating these empirically-observed dengue cycles into our system improved our ability to predict non-epidemic years by approximately 20% (specificity=69%) and increased overall accuracy to 74.2% (Table 1). Specifically, the additional decision rule replaced 7 epidemic forecasts with non-epidemic forecasts, of which 5 were correct (Fig. 3B). A majority of these cases belonged to cities which had experienced 3 consecutive epidemic years leading up to the prediction.

Overall, the combined approach (weather-based plus dengue cycles) was dominantly driven by weather patterns, and informed by the decision rule only in a few cases when historical data showed a very strong likelihood of either an epidemic or not epidemic year to happen. Thus, the decision rule to favor the Markov model acts as an “expert opinion” for situations in which there is clear evidence that a given predicted scenario (even if suggested by the weather patterns) is unlikely. Our specific finding - that the dengue cycles were used exclusively to overturn epidemic forecasts - suggests that while the weather conditions in those locations and years were identified to be conducive to an outbreak, there was stronger evidence that the population may have had low susceptibility to infection (thus avoiding an outbreak), based on multiple consecutive preceding years of high disease incidence.

### Model performance by year

The success of our combined epidemic forecasts varied by year, reflecting the difficulty of forecasting disease activity relying only on weather patterns and the empirically extracted susceptibility cycles. During the last three years of the time series (2015-2017), epidemics were predicted by the weather-only models with at least 80% accuracy, with 100% of the 13 outbreaks in 2016 correctly forecasted (Fig. 3B,C). Conversely, non-epidemic years during 2013-2014 were particularly difficult to predict, with only one-third and one-half of cities correctly forecasting non-epidemics for these years, respectively. The most successful non-epidemic predictions occurred in 2012, for which 6 out of 8 non-epidemics (75%) were predicted correctly. Overall, 2015 and 2016 were the most successfully classified years, with 80% and 85% of municipalities correctly classified as epidemics or non-epidemics, respectively, while 2014 and 2017 were the most difficult years to predict, with 45% and 35% of municipalities misclassified, respectively.

Incorporating information on the dengue cycles helped detect an additional non-epidemic in 2012 and 2015, and an additional 3 non-epidemics in 2017 (Fig. 3B).

### Quantifying the strength of predictions

Because our forecast system produces deterministic binary predictions (epidemic/non-epidemic year) using local-in-time support vector machine classifiers, a natural question is how to quantify the conviction (or confidence) of each prediction. It is important to note that the number of observations per city is small (n=17), and thus, a rigorous probabilistic approach to quantify conditional probabilities of success is not feasible. However, in the interest of better communicating to public health officials the reliability of our predictions in a given location and time period, as well as to identify the determinants of success of our prediction system if one were to extend our predictive approach to new locations, we explored simple ways to characterize the accuracy and conviction of predictions. We did this based on both the historic performance of the selected ensemble generating the prediction, as well as the performance of the weather-based classifiers themselves.

Our prediction system combines the output of a collection of local-in-time binary classifiers that use different time periods (characterized by an initial point in time, t0, and a window length, p), prior to the typical date of the onset of dengue outbreaks, as predictors. For each city and each year, the combination of these outputs is calculated using a voting system that only considers time windows that have consistently exhibited the highest historical out-of-sample prediction performance among all other time windows of the calendar year. In our framework, time windows are automatically selected into the forecasting ensemble if (a) their own historic out-of-sample performance is high and (b) the historical performance of their calendar neighbors, that is, models using temporally nearby time windows as predictors, is high as well.

Consequently, we computed metrics of ensemble accuracy and strength (or confidence) by quantifying both of these elements. We found that in cities where the predictive performance of our approach is highest (Fig. S2), the successful individual classifiers that contribute to our final prediction use as input temporal regions that are clustered around one another (as shown in Fig. 4), suggesting that the presence of temporally consistent weather patterns can be thought of as an indicator of success of our methodology.

**Figure 4.**
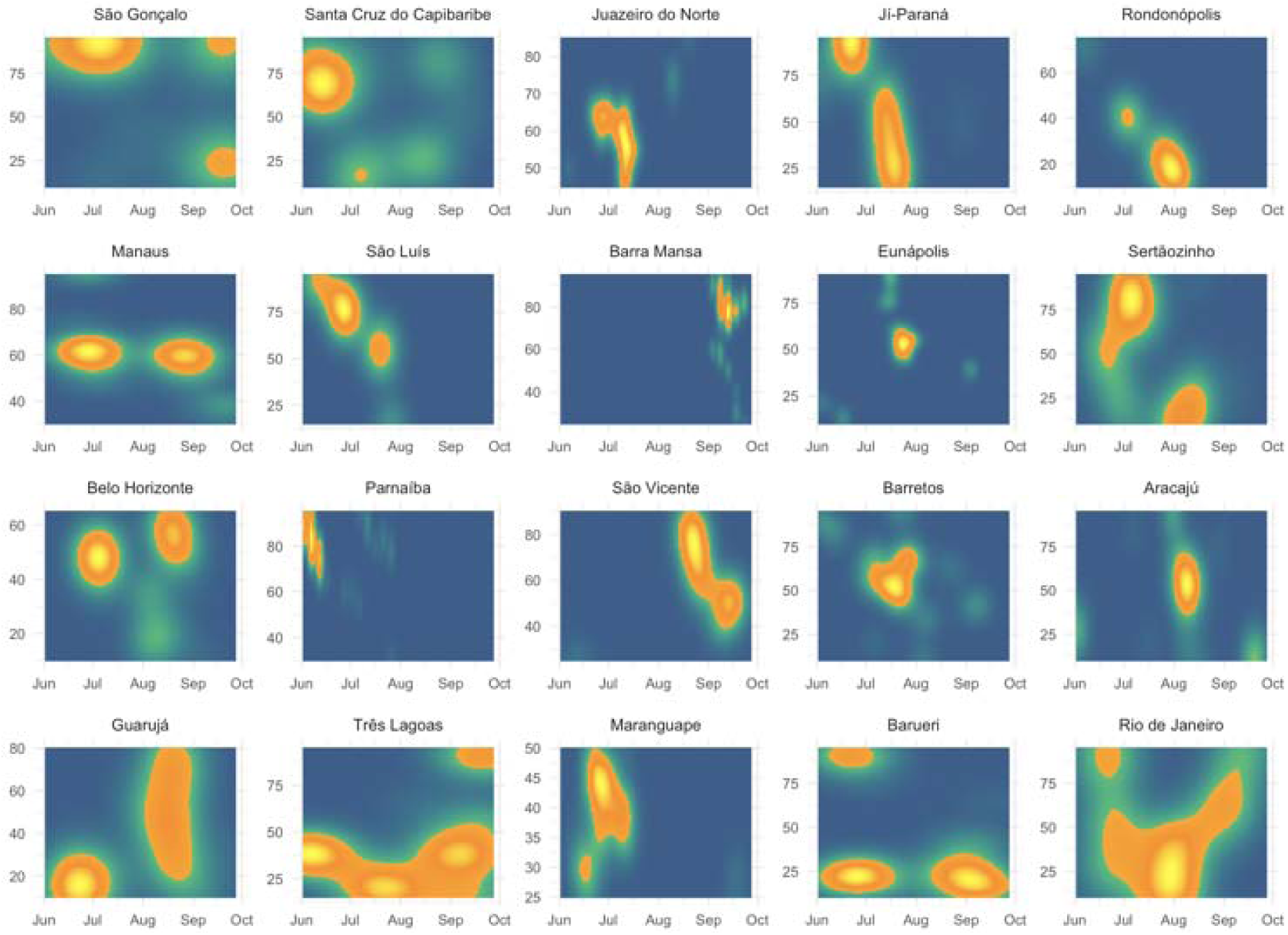
Periods of the year selected into the ensemble forecast model for 2012-2017, by municipality. The x-axis (t_0_) indicates the start date of the time interval, and the y-axis (p) indicates the length of the time interval from which weather data were gathered (10-95 days).Municipalities with smaller and brighter yellow centers are those which exhibit the highest consistency in the predictive performance of weather patterns. Municipalities are ordered by decreasing ensemble prediction accuracy; that is, the proportion of years correctly forecasted by the ensemble method over the years 2012-2017.

It is important to note that models with high historic prediction performance may still lead to poor outcomes if the weather data for the year of (out-of-sample) forecast does not clearly belong to an epidemic or non-epidemic class, as learned by the individual classifiers, and/or if its weather patterns happen to “look like” those appearing historically in the opposite class.

In order to further assess the individual strength or conviction of each individual classifier, we estimated whether the separability or difference between the two classes (epidemic vs non-epidemic) was well captured by the classifier by extracting calibrated posterior probabilities of each SVM model using Platt’s scaling^33^. The posterior probability reflects the distance to the separation boundary distinguishing epidemic and non-epidemic years on the basis of weather. Thus, a higher probability represents how strongly the weather patterns of the prediction year aligned with those experienced by prior outbreak or non-outbreak years. We observed that in general, the probabilities were moderately calibrated, i.e. roughly 80% of predictions made with 0.8 probability were epidemics (Fig. S3); however, the small sample size (i.e. six out-of-sample years for each of the 20 cities) limits the ability to interpret this feature appropriately. We found that this measure of separability was not a particularly good indicator of accuracy; that is, our approach failed even in scenarios with high separability. Several factors may be driving this finding, including insufficient training data and the influence of factors beyond weather (e.g. sociodemographic characteristics, land use) on outbreaks; we elaborate further in *Supplementary Material*.

Both approaches to characterize the confidence of our predictions – quantifying ensemble strength and quantifying the separability of the data – highlight separate limitations of our modeling framework. First, we expect that both a greater variety of environmental variables (e.g. humidity, vegetation, standing water) and non-environmental variables (e.g. human activity and public health interventions) will contribute to more accurate predictions by considering broader factors that contribute to dengue fever activity in a given location. Second, the robustness of our predictions was limited by a short time series of annual information, which may not be sufficient to detect clear differences in epidemic and non-epidemic years on the basis of weather alone. Nonetheless, our reproducible modeling framework can easily be extended to accommodate additional predictors and longer time series, and thus we highlight these as limitations of only the present case study, with potential for improved performance in other data settings.

## Discussion

Here we have presented a novel approach to forecast dengue fever outbreak years in Brazil at its smallest administrative unit, the city level, using a single, dynamic and flexible modeling framework that uses only two weather variables and historical information on yearly dengue activity. Our approach automatically learns from weather and population susceptibility patterns of any inputted yearly time series of dengue incidence and leverages the best historic predictions to generate an ensemble forecast. We find that complementing our weather-based statistical approach with observed 3-4 year cycles of dengue fever outbreaks (as a proxy for population susceptibility) is key for our models to achieve higher accuracy and improve substantially in predicting non-epidemic years. These forecasts may provide timely information on dengue fever activity to policymakers months ahead of outbreak seasons. Further, our entirely data-driven models show an ability to learn from complex relationships between dengue epidemics and climatic conditions and identify, in vastly different locations, potentially relevant weather patterns with likely biological significance. Importantly, these models can be immediately extended to other locations, requiring no location-specific manipulation or inputs aside from a globally-available time series of daily temperature and precipitation as well as a complete yearly record of Dengue incidence.

Using weather information only, our models seek to characterize and exploit the predictive ability of distinct weather patterns preceding outbreak years. Because our framework automatically identifies the time periods for which weather patterns produce strong signals, it was possible to identify temporal weather signatures in multiple locations with vastly different ecosystems and geographic locations. For this, we observed that cities with better overall prediction accuracy had stronger weather signatures, suggesting perhaps some biological consistency. For example, the southeastern municipality of Barra Mansa (5 of 6 ensemble years predicted correctly) exhibited strong signals from time windows spanning the first half of the city’s rainy season, in October through December of each year. Farther north, the hot, wet, and humid municipality of Manaus (5 of 6 ensemble years predicted correctly), situated at the mouth of the Amazon, appeared to show two distinct weather signatures straddling the driest month of the year, August. These patterns, generated from 10 years of out-of-sample model predictions, suggest that in different regions of Brazil, weather may affect dengue transmission differently and at different times of the year. However, in locations where weather-based predictions were less successful, these signatures were not distinct; for instance, Rio de Janeiro (3 out of 6 ensemble years predicted correctly) showed no clear temporal trend. In cities such as these, we might expect to see a lower influence of weather patterns on transmission compared to other predictors (e.g. policy, population behavior, human land use). We did not find clear patterns by geography, population density, or municipality size. We believe this work should catalyze important research both on the local influence of weather patterns on dengue outbreaks as well as the extent to which other factors drive outbreaks in these locations. Moreover, this data-driven approach may help generate hypotheses on the relevance of multiple factors that may influence the dynamics of seasonal dengue outbreaks.

Even weather conditions that appear highly suitable for an outbreak (or none), based on historical information, may be challenged by other factors that limit (or encourage) transmission of dengue. A key strength of our approach is the incorporation of empirically-observed information on dengue fever susceptibility cycles, to correct for potential short-term immunity that results from previous exposure to the dengue virus. We found that these susceptibility cycles were critical to the performance of models, particularly those which identified weather patterns suitable for a dengue outbreak in a year with potentially low population susceptibility to infection. For instance, this approach correctly identified 3 additional non-epidemics in 2017 compared to weather patterns alone, supporting the discourse on the unusually low dengue activity seen in Brazil in 2017^34^. Still, our models missed half (6/12) of non-epidemics in 2014, which was predicted by experts to be a low transmission year due to immunity provided by a large 2013 outbreak with no changes in circulating DENV serotypes^35,36^. Thus, incorporating information on specific circulating serotypes could be used to better detect changes in population immunity and enhance our approach. Empirical and modeling-based seroprevalence studies may aid with this component, though this surveillance information is more challenging to routinely acquire^37^. Regardless, here we highlight the importance of incorporating mechanistic processes of disease transmission into data-driven approaches that may be otherwise blinded to them.

Because dengue transmission is driven by multiple complex socioecological and biological factors, we expect our models to capture only a portion of the epidemiologic triangle. Here we show the performance of two simple and relevant weather indicators of dengue fever, but the incorporation of additional weather features (i.e. humidity, vegetation, soil water absorption) combined with a feature selection step may lead to improved accuracy of forecasts, by considering more complex weather conditions preceding dengue outbreaks. Still, we demonstrate robustness of our approach by replicating the study using an alternative feature extraction method, singular value decomposition, with similar results (Fig. S4, *Supplementary Materials & Methods*). Further, weather- and susceptibility-based models can contribute valuable information to larger ensemble approaches leveraging a collection of mobility, sociodemographic, epidemiologic, climatic, and biological information.

Our approach also demonstrates the feasibility (and limitations) of predicting in a “small data” setting, wherein only 17 outcome data points were available in total for training and out-of-sample predictions (each representing annual outbreak status between 2001-2017). We chose a short training period (initial 7 years) to maximize the number of out-of-sample predictions, but ultimately it is difficult to establish strong climatic distinctions between outbreak and non-outbreak years in the data with so few samples. Thus, we anticipate improvement in performance for settings that have multiple decades of data, which would allow for longer training periods, improved separability in the data, and more stable identification of dengue susceptibility cycles, all improving the quality and robustness of predictions.

Ultimately, this framework provides a simple, reproducible method of predicting dengue fever outbreak years in a wide range of locations. Given that the global and economic burden of dengue is placed at an estimated 390 million infections and $8.9 billion per year^12,38^, optimizing resource allocation for the disease prevention is critical. However, control of the *Aedes* mosquito requires weeks or months before effects are seen on the vector population, so predicting dengue outbreaks up to several months of their onset is ideal. Our reproducible approach, which uses of globally-available data with daily resolution, is intended to serve as an unsupervised learning framework to produce early outbreak warnings in any desired context, resulting in more efficient resource mobilization, budgeting, and prevention campaigns. Developing transparent early warning systems at the local level is emerging as a top global health priority, making our contribution both timely and impactful.

## Materials and Methods

We developed a single, flexible modeling framework capable of identifying potentially useful weather patterns to predict dengue fever, and used this to forecast annual outbreak status (epidemic / non-epidemic).

Our workflow, outlined in Fig. 1, combines elements from signal processing/spectral analysis, machine learning, and ensemble modeling to achieve robust, data-driven epidemic forecasts that do not require any prior knowledge of the system (i.e. climatic influences on dengue transmission). Our research question is inherently one of time series classification, to forecast epidemic vs. non-epidemic years of dengue fever. The workflow begins with a time series of hourly and daily weather information, which serve as inputs to a collection of classifiers that contribute to ensemble-based epidemic predictions. Our approach can be described in 5 steps:

1. *Signal preprocessing*: for a time series of weather data, define time intervals of varying sizes (10-95 days across the last 7 months of the calendar year), and use a windowing technique^31^ to include information within several days of the interval. In contrast to^31^, there are no deleterious effects due to missing temperature data since it is acquired via satellite instead of ground measurements.
2. *Time series feature extraction*: extract a simple summary measure for 2 weather variables with known influence on mosquito-borne disease dynamics, temperature and frequency of precipitation. Although more variables can be considered, they have little influence on the predictive power in comparison with the two selected^31^.
3. *Independent model training and prediction*: train a collection of independent support vector machine (SVM) classifiers on historical information from each unique time interval, and generate an out-of-sample epidemic prediction for the following year. Although SVM was used in^31^, we provide here a richer out-of-sample prediction scheme for forecasting.
4. *Model selection*: choose the best 11 models, representing strongly predictive periods of the year preceding outbreaks, based on a) historical out-of-sample prediction accuracy and b) out-of-sample performance of neighboring time intervals
5. *Ensemble prediction*: determine a final out-of-sample epidemic forecast by majority vote of the selected top models

To potentially enhance the performance of this exclusively weather-based approach, we implemented a post-hoc step incorporating empirical information on 3- and 4-year dengue fever cycles as a proxy for population susceptibility to infection.

1. *Dengue cycles*: implement a decision rule governed by the second- and third-order Markov transition probabilities, reflecting the transition between consecutive sequences of epidemic and non-epidemic states

We applied our approach to 20 cities in Brazil spanning large geographic and population ranges (Fig. S1, Table S1). We used as input a historical time series spanning 17 years and consisting of information on dengue case reports (number, annual) and 2 weather variables: 2-meter air temperature (Kelvin, daily) and precipitation (kg/m^2^, hourly). We describe data sources, acquisition, and processing in the *Supplementary Materials*. After an initial training period of 7 years, we generated 10 years of out-of-sample epidemic predictions for each of the independent models using a one-year expanding training window (Step 2). We used the first 4 years of out-of-sample predictions to inform ensemble model selection (Step 4), and produced ensemble-based predictions for the remaining 6 years (Step 5).

### Signal preprocessing

Using a daily time series of weather data to forecast dengue fever epidemic status requires identifying the most predictive period(s) of the calendar year during which weather information contains a strong signal for subsequent dengue fever outbreaks. In order to construct a single framework that can automatically identify important weather signals in multiple different locations with vastly different ecosystems and weather patterns, we allow the data to inform the choice of time intervals. Our algorithm achieves this by scanning over multiple, partially-overlapping time intervals across the calendar year, and building hundreds of models on these different intervals in order to select those with the strongest signals.

Each time interval is defined by a start date, *t0*, between early June and late September, and a period length, *p*, of between 10 and 95 days. The combination of each (*t0, p*) produces multiple, partially-overlapping intervals spanning the last 7 months of the calendar year.

Borrowing from spectral analysis and wavelet decomposition, we use a windowing-inspired approach to better capture signals within the time intervals. Windowing is typically used to improve signal clarity, and here we apply a rectangular “range” as described in^31^ to incorporate information in the days both within and around each time interval. We define a rectangle of 5 × 6, indicating that, for every defined (*t0, p*) time interval, the algorithm collects information from 5 consecutive start dates, *t0, t0+1, …, t0+4*, spanning 6 consecutive period lengths, *p, p+1*, …, *p+5*. Each time interval and weather variable, then, is summarized by 30 data points, each capturing slightly different temporal slices from the time series. This process effectively adds a bit of redundant information to the model building process - to which our learning algorithm, the support vector machine, is in general robust - in order to pick up signals in the data that may not be captured by applying an arbitrary “start” and “end” cutoff to the data.

### Time series feature extraction

Time series data must be transformed into appropriate inputs in order to be used in supervised learning models. This process, called time series feature extraction, involves computing summary features of the time series, which can range from simple means to complex wavelet transforms. To test the feasibility of our approach using only simple summary features, we extracted the following features within each (*t0, p*) time interval based on the findings of^31^: 1) the arithmetic mean of daily temperature, and 2) mean precipitation frequency, with frequency defined as the time interval (in days) between peaks (local maxima) of daily precipitation. In the *Supplementary Materials*, we present an alternative method of feature extraction using singular value decomposition.

### Independent model training and prediction

The goal of our independent model building step is to identify dynamically, through the continually-updating performance of a collection of models, the periods of the year that are most predictive of annual dengue outbreaks, in order to exploit a small number of them to generate forecasts.

To forecast outbreak years, we trained a collection of support vector machine (SVM) classifiers on an initial 7 year training period, and produced annual forecasts incorporating the most recently available weather information using a dynamic, one-year expanding training window. A unique SVM was trained for each of the (*t0, p*) time intervals, resulting in a total of 432 independent models trained per year. Each model generated out-of-sample predictions for the remaining 10 years of data. Predictions were made by classifying the 30 out-of-sample data points corresponding to the weather information preceding the target year, and taking a majority vote. In order to handle highly nonlinear relationships between weather variables, both radial basis function (RBF) and sigmoid kernels were used and evaluated for performance, and show results for the best respective kernel in each city. We tuned model parameters (gamma, soft margin cost function, and coefficient) using 10-fold cross-validation.

Support vector machines, a supervised learning method for classification, were used because of their flexibility in the face of complex, nonlinear decision boundaries and their robustness to overfitting and outliers. The property that underpins these advantages is known as the “large-margin classifier.” SVMs are also known for their good performance in high-dimensional feature space, which is advantageous for the scale-up of the model to include dozens more predictors.

### Model selection

From the resulting collection of 432 models, the best-performing models (n=11) were selected each year based on a) historical out-of-sample prediction accuracy (% outbreak forecasts correct) and b) out-of-sample prediction accuracy of neighboring models (representing similar time intervals). These models thus represent strongly predictive periods of the year preceding outbreaks, and the algorithm rewards the high performance of similar temporal windows over the high performance of a time window whose neighbors exhibit poor prediction tendencies. Because the model building process is dynamic, resulting in a new collection of models each year with continually-updating performance measures, the selection of the 11 models changes from year to year.

In order to get a sense of the out-of-sample performance of the 432 models, we allowed all models to generate 4 years of out-of-sample predictions before the top 11 models were selected based on this prediction accuracy. As a result, the ensemble approach, which exploited the predictions of the top 11 models, was used for the final 6 years of out-of-sample predictions.

### Ensemble prediction

Ensemble learning helps improve machine learning algorithms by combining the results of multiple trained predictors in order to generate a single, robust prediction. In our approach, we combine the results from the strongest-performing models, which represent the most highly predictive time periods preceding dengue outbreaks. While there are an abundance of ensembling methods in machine learning, we use a simple majority vote of the 11 models to decide a single forecast. These single forecasts were produced for the last 6 years of the 17-year dataset, representing the culmination of a prediction process that involves: 7-year initial training period, 4-year out-of-sample model calibration period, and 6-year out-of-sample ensemble prediction period. Across 20 Brazilian municipalities, this scheme produced 120 municipality-years of out-of-sample ensemble predictions.

### Dengue cycles

Our weather-based ensemble approach remains ignorant to the relationship between weather patterns and dengue outbreaks, instead allowing the data to drive model selection and predictions. However, endemic transmission of dengue fever is typically distinguished by periodic outbreak cycles of around 3-4 years. These outbreak cycles are thought to occur as a result of 1) an exhaustion of susceptibles after an outbreak, and 2) and short-term cross-immunity to other circulating DENV serotypes after infection^21^. Both factors result in a depletion of the population vulnerable to infection, and act as barriers to subsequent outbreaks. Independent of climate variability over the years, we expect some preservation of these cycles.

Consequently, we implemented a “decision rule” in the model based on the observed transitions between epidemic- and non-epidemic years across 51 Brazilian municipalities meeting endemic inclusion criteria (Supplemental Information). Across these municipalities, we computed the mean second- and third-order Markov transition probabilities, representing the probability of transition from one outbreak state (epidemic/non-epidemic) to the opposite outbreak state (non-epidemic/epidemic) after 2 and 3 consecutive years, respectively. Thus, we obtained the transition probabilities governing the following 3- and 4-year cycles: 001, 110, 0001, and 1110 (0= non-epidemic year, 1= epidemic year). Transition probabilities were computed based only on the first 11 years of data; that is, the years preceding the 6 out-of-sample ensemble predictions.

Our decision rule acts as a surrogate “expert opinion,” overturning the ensemble prediction if the probability of a specific transition exceeded the percent of model votes (out of 11 votes). For example, if the ensemble predicts an epidemic year to succeed 2 epidemic years with 7 votes, the corresponding “strength” of that vote is 63% (7/11), which is weaker than the corresponding observed second-order transition probability for a non-epidemic year to follow 2 epidemic years (0.71). In this case, the model vote would be overridden to predict a non-epidemic year instead of an epidemic year.

We compared the performance of predictions based solely on weather patterns to those which incorporate additional empirical data from outbreak cycles.

## Data Availability

The epidemiologic data used in this study are available from the Brazilian Ministry of Health DataSUS: http://www2.datasus.gov.br/DATASUS/index.php?area=0203&id=29878153. Meterological data (MERRA-2) are available through the Global Modeling and Assimilation Office (GMAO) at NASA Goddard Space Flight Center: https://disc.gsfc.nasa.gov/datasets?keywords=%22MERRA-2%22&page=1&source=Models%2FAnalyses%20MERRA-2.

## Author contributions

S.F.M., J.N.K., and M.S. conceived the study; S.F.M., J.N.K., and M.S. formulated the experimental design; S.F.M. collected the data; S.F.M. C.L.C. analyzed the data; and all authors discussed results, contributed to manuscript preparation, and reviewed the manuscript.

## References

1. Ford, T. E. et al. Using Satellite Images of Environmental Changes to Predict Infectious Disease Outbreaks. Emerg. Infect. Dis. 15, 1341–1346 (2009).

2. Sewe, M. O., Tozan, Y., Ahlm, C. & Rocklöv, J. Using remote sensing environmental data to forecast malaria incidence at a rural district hospital in Western Kenya. Sci. Rep. 7, 2589 (2017).

3. McGough, S. F., Brownstein, J. S., Hawkins, J. B. & Santillana, M. Forecasting Zika Incidence in the 2016 Latin America Outbreak Combining Traditional Disease Surveillance with Search, Social Media, and News Report Data. PLoS Negl. Trop. Dis. 11, e0005295 (2017).

4. Yang, S., Santillana, M. & Kou, S. C. Accurate estimation of influenza epidemics using Google search data via ARGO. Proc. Natl. Acad. Sci. U. S. A. 112, 14473–14478 (2015).

5. de Almeida Marques-Toledo, C. et al. Dengue prediction by the web: Tweets are a useful tool for estimating and forecasting Dengue at country and city level. PLoS Negl. Trop. Dis. 11, e0005729 (2017).

6. Bengtsson, L. et al. Using mobile phone data to predict the spatial spread of cholera. Sci. Rep. 5, 8923 (2015).

7. Kramer, A. M. et al. Spatial spread of the West Africa Ebola epidemic. R Soc Open Sci 3, 160294 (2016).

8. Zhu, Z. et al. Comparative genomic analysis of pre-epidemic and epidemic Zika virus strains for virological factors potentially associated with the rapidly expanding epidemic. Emerg. Microbes Infect. 5, e22 (2016).

9. Dudas, G. et al. Virus genomes reveal factors that spread and sustained the Ebola epidemic. Nature 544, 309–315 (2017).

10. Reich, N. G. et al. A collaborative multiyear, multimodel assessment of seasonal influenza forecasting in the United States. Proc. Natl. Acad. Sci. U. S. A. 116, 3146–3154 (2019).

11. Buczak, A. L. et al. Ensemble method for dengue prediction. PLoS One 13, e0189988 (2018).

12. Bhatt, S. et al. The global distribution and burden of dengue. Nature 496, 504–507 (2013).

13. Stanaway, J. D. et al. The global burden of dengue: an analysis from the Global Burden of Disease Study 2013. Lancet Infect. Dis. 16, 712–723 (2016).

14. Morin, C. W., Comrie, A. C. & Ernst, K. Climate and dengue transmission: evidence and implications. Environ. Health Perspect. 121, 1264–1272 (2013).

15. Tjaden, N. B., Thomas, S. M., Fischer, D. & Beierkuhnlein, C. Extrinsic Incubation Period of Dengue: Knowledge, Backlog, and Applications of Temperature Dependence. PLoS Negl. Trop. Dis. 7, e2207 (2013).

16. Rohani, A., Wong, Y. C., Zamre, I., Lee, H. L. & Zurainee, M. N. The effect of extrinsic incubation temperature on development of dengue serotype 2 and 4 viruses in Aedes aegypti (L.). Southeast Asian J. Trop. Med. Public Health 40, 942–950 (2009).

17. Liu, Z. et al. Temperature Increase Enhances Aedes albopictus Competence to Transmit Dengue Virus. Front. Microbiol. 8, (2017).

18. Byttebier, B., De Majo, M. S. & Fischer, S. Hatching Response of Aedes aegypti (Diptera: Culicidae) Eggs at Low Temperatures: Effects of Hatching Media and Storage Conditions. J. Med. Entomol. 51, 97–103 (2014).

19. Barry W. Alto, D. B. Temperature and Dengue Virus Infection in Mosquitoes: Independent Effects on the Immature and Adult Stages. Am. J. Trop. Med. Hyg. 88, 497 (2013).

20. Scott, T. W. et al. Longitudinal studies of Aedes aegypti (Diptera: Culicidae) in Thailand and Puerto Rico: population dynamics. J. Med. Entomol. 37, 77–88 (2000).

21. Adams, B. et al. Cross-protective immunity can account for the alternating epidemic pattern of dengue virus serotypes circulating in Bangkok. Proc. Natl. Acad. Sci. U. S. A. 103, 14234–14239 (2006).

22. Mbogo, C. M. et al. Spatial and temporal heterogeneity of Anopheles mosquitoes and Plasmodium falciparum transmission along the Kenyan coast. Am. J. Trop. Med. Hyg. 68, 734–742 (2003).

23. Acevedo, M. A. et al. Spatial heterogeneity, host movement and mosquito-borne disease transmission. PLoS One 10, e0127552 (2015).

24. Torres-Sorando, L. & Rodríguez, D. J. Models of spatio-temporal dynamics in malaria. Ecol. Modell. 104, 231–240 (1997).

25. Teurlai, M. et al. Socio-economic and Climate Factors Associated with Dengue Fever Spatial Heterogeneity: A Worked Example in New Caledonia. PLoS Negl. Trop. Dis. 9, e0004211 (2015).

26. Descloux, E. et al. Climate-based models for understanding and forecasting dengue epidemics. PLoS Negl. Trop. Dis. 6, e1470 (2012).

27. Guo, P. et al. Developing a dengue forecast model using machine learning: A case study in China. PLoS Negl. Trop. Dis. 11, e0005973 (2017).

28. Chuang, T.-W., Chaves, L. F. & Chen, P.-J. Effects of local and regional climatic fluctuations on dengue outbreaks in southern Taiwan. PLoS One 12, e0178698 (2017).

29. Johansson, M. A., Reich, N. G., Hota, A., Brownstein, J. S. & Santillana, M. Evaluating the performance of infectious disease forecasts: A comparison of climate-driven and seasonal dengue forecasts for Mexico. Sci. Rep. 6, 33707 (2016).

30. Lauer, S. A. et al. Prospective forecasts of annual dengue hemorrhagic fever incidence in Thailand, 2010–2014. Proceedings of the National Academy of Sciences 115, E2175–E2182 (2018).

31. Stolerman, L., Maia, P. & Kutz, J. N. Data-Driven Forecast of Dengue Outbreaks in Brazil: A Critical Assessment of Climate Conditions for Different Capitals. (2016).

32. Guzman, M. G., Alvarez, M. & Halstead, S. B. Secondary infection as a risk factor for dengue hemorrhagic fever/dengue shock syndrome: an historical perspective and role of antibody-dependent enhancement of infection. Arch. Virol. 158, 1445–1459 (2013).

33. Platt, J. Probabilistic Outputs for Support Vector Machines and Comparisons to Regularized Likelihood Methods. Advances in large margin classifiers 10, 61–74 (1999).

34. Lopes, T. R. R., Silva, C. S., Pastor, A. F., Silva, J. V. J. & Júnior. Dengue in Brazil in 2017: what happened? Rev. Inst. Med. Trop. Sao Paulo 60, (2018).

35. van Panhuis, W. G. et al. Risk of Dengue for Tourists and Teams during the World Cup 2014 in Brazil. PLoS Negl. Trop. Dis. 8, e3063 (2014).

36. Massad, E. et al. Risk of symptomatic dengue for foreign visitors to the 2014 FIFA World Cup in Brazil. Mem. Inst. Oswaldo Cruz 109, 394–397 (2014).

37. Honório, N. A. et al. Spatial Evaluation and Modeling of Dengue Seroprevalence and Vector Density in Rio de Janeiro, Brazil. PLoS Negl. Trop. Dis. 3, e545 (2009).

38. Shepard, D. S., Undurraga, E. A., Halasa, Y. A. & Stanaway, J. D. The global economic burden of dengue: a systematic analysis. Lancet Infect. Dis. 16, 935–941 (2016).

39. Gelaro, R. et al. The Modern-Era Retrospective Analysis for Research and Applications, Version 2 (MERRA-2). J. Clim. 30, 5419–5454 (2017).

40. Hendrycks, D. & Gimpel, K. A Baseline for Detecting Misclassified and Out-of-Distribution Examples in Neural Networks. arXiv: 1610.02136v3 (2018).

