## Supplementary Materials for "Combining weather patterns and cycles of population susceptibility to forecast dengue fever epidemic years in Brazil: a dynamic, ensemble learning approach"

***All authors have read/approved the manuscript. This manuscript has not been published elsewhere. No writing assistance was provided in preparation. Data used in the study are from openly available sources and are described/referenced accordingly.***

**Contents of Supplementary Material:**

Supplemental Materials & Methods

Supplemental Results

Table S1

Figs. S1-S7

Supplemental Materials & Methods

**Data**

*Study sites*

We tested our model on a time series of 17 years (2001-2017) for 20 municipalities in Brazil meeting the following criteria: 1) having experienced between 7 and 10 epidemic years during the 17-year time period, with no more than 70% of epidemic years occurring in either half of the time period; and 2) having a population over 100,000 by 2017. The first criterion captures a loose definition of a “dengue-endemic” location, which is convenient in two ways: first, locations which experience only epidemic or only non-epidemic years are likely experiencing disease dynamics unrelated to annual changes in weather patterns, and are thus not appropriate for our model; and second, it ensures that our model is able to train initially on both classes (epidemic and non-epidemic year) for each location before making out-of-sample predictions. In accordance with Brazilian Ministry of Health standards, we defined an epidemic year to be a year in which the number of confirmed cases of dengue fever exceeds 100 per 100,000 persons.

The municipalities included in the study span a wide geographic range (14 Brazilian states) and range in land area from 24 to 4000 mi^2^, in starting population (population in 2001) of between 87,000 and 6 million, and in starting population density from 24 to 13,000 persons/mi^2^ (Table S1).

*Epidemiologic data*

The number of confirmed cases of dengue fever are reported annually at the municipal level and made publicly available from the Brazilian Notifiable Disease System (SINAN) for the years 2001-2012. We obtained weekly case numbers at the municipal level from the Brazilian MOH for the years 2013-2015, and from local municipal governments and epidemiologic reports for 2016 and 2017.

*Demographic data*

Population estimates by year and municipality were obtained as publicly-available data from the Brazilian Institute of Geography and Statistics (IBGE).

*Weather data*

Globally modeled and assimilated weather data were obtained from the Modern Era Retrospective-analysis for Research and Applications, Version 2 (MERRA-2)^39^. The MERRA-2 data are publicly available through the Global Modeling and Assimilation Office (GMAO) at NASA Goddard Space Flight Center. We obtained daily temperature at 2 meters (mean, K) and hourly precipitation (kg/m^2^) at a native grid resolution of 0.5° x 0.625° and extracted these to municipalities by overlaying a spatial file of municipality boundaries and taking the weighted average of the grid cells covering municipal boundaries. We calculated the total accumulated rainfall in a day (mm) as the sum of hourly precipitation (kg/m^2^/hr, which is equivalent to mm/hr) over the 24-hour period. We show the time series of mean daily temperature (K) and total precipitation (mm) in Fig. S5.

**Ensemble strength calculations**

We computed a simple metric to quantify the strength of the 11-model ensemble used to make yearly forecasts for each municipality. For each model, the metric was the sum of (a) the historical out-of-sample forecast accuracy of the time window, computed as the number of years correctly predicted divided by N prior years, and (b) the average historical out-of-sample accuracy of the time window and its (up to) 8 surrounding neighbors (t_0_ ± 5, p ± 5). Thus, the maximum possible strength for any given ensemble was 2.0 (1.0 + 1.0).

**Alternative ensemble method**

To assess whether our feature extraction methodology was comprehensively extracting meaningful weather patterns predictive of dengue epidemic years, we implemented an additional feature extraction methodology based on the Singular Value Matrix Decomposition (SVD) as introduced in^31^. In this alternative approach, a pair (t0, p) denotes a single time interval, during a given year, specified by the pair of values assigned to t0 (day of the year) and p (length of the time interval). For a given (t0, p), our approach extracts all the weather data associated to the identified time window throughout the years in our data set. Subsequently, a classifier aimed at predicting whether a subsequent year will be epidemic was constructed using as input the set of historically observed singular values associated to the dominant mode of the SVD, on each of the (t0, p), for a dynamically increasing training set. For historically successful classifiers associated to a given (t0,p), a voting system was designed to produce a final estimation of the likelihood of a year to be epidemic. More details are presented below.

*Signal preprocessing*

For a single pair (t0, p) and N years, we extracted the observed weather variables’ time-series within the (t0, p) time interval for each year. We then normalized the data by calculating the temporal z-score for both meteorological variables. Then, we built a 2p by N matrix A. Each column of this matrix consists of the two meteorological variables concatenated (dimension 2p) for a given year, totaling N columns (years).

*Feature extraction*

For a given training number of years, we obtained the Singular Value Decomposition (SVD) of the associated matrix A. The SVD operation decomposes a matrix A into a product of matrices USV* that can be used to find a more compact representation of the data matrix A by projecting every column vector of our matrix into a smaller space spanned by a subset of the columns of U, which can be seen as an orthogonal basis generated by the SVD. The SVD “compresses” the matrix into a linear combination of columns of U. The largest singular value (contained in the matrix S) is associated to the column that captures the maximum variance of the column space of A and contains most of the information of A. Further details can be found in^31^.

*Independent model training and prediction*

To produce an out-of-sample prediction for a year on a single (t0, p) in our alternative approach, we included all available data prior to that year, applied the pre-processing step, then separated the data into a training dataset comprised of data from the first year available up to 2 years before the prediction year. The data from 1 year before the prediction was then used as our out-of-sample input to our ensemble in ends to classify next year as either epidemic or not epidemic. We applied the SVD on the training data and fit a classifier on the compressed version of A. Finally, we transformed our out-of-sample input into the new representation created by the SVD and used the classifier to detect if the prediction year would be classified as having an outbreak or not. For our classification method, we decided to fit an SVM using a non-linear kernel (Radial Basis Function) to our training dataset, allowing for more complex decision boundaries when separating our training data. Using the same domain for the variables t0 and p described in the main approach, and given we do not use the windowing method in this alternative approach, we had a total of 12960 possible classifiers for the June to late-September period of prediction for every year.

*Model selection*

For all 12960 possible (t0, p), we selected the best performing models based on historical out-of-sample prediction accuracy and an accuracy threshold that was set -20% on the highest recorded accuracy percent. In contrast to ensemble that utilizes the windowing approach, we let our accuracy threshold decide the number of models to vote in the ensemble.

Historical out of sample accuracy was based on the performance of predicting an out-of-sample year six years prior to our out-of-sample prediction. The out-of-sample accuracy for a specific (t0, p) was constructed as the result of the ratio between the number of years where the classifier successfully predicted the out-of-sample year and the total number of years in this window.

*Ensemble approach*

For an out-of-sample prediction, we select a subset of all possible combinations of (t0, p) as the classifiers that will participate in a simple voting system. We then fit a classifier in this selected group of (t0, p) combinations and sum each decision (assigning the outbreak label a +1 and a non-outbreak label a -1) weighted by its historical out-of-sample accuracy to avoid any possible draws and produce a final decision for the out-of-sample year. This alternative approach also produced forecasts for the last 6 years of the 17-year dataset.

Supplemental Results

**Quantifying separability of the data used for predictions**

We estimated whether the separability or difference between the two classes (epidemic vs non-epidemic) was well captured by the classifier by extracting calibrated posterior probabilities of each SVM model using Platt’s scaling. The posterior probability reflects the distance to the separation boundary distinguishing epidemic and non-epidemic years on the basis of weather. Thus, a higher probability represents how strongly the weather patterns of the prediction year aligned with those experienced by prior outbreak or non-outbreak years.

Across the models composing each ensemble SVM, we computed the mean of the posterior probabilities (MPP, Fig. 3A) and compared these to the accuracy of the prediction. We found that a majority (75%) of municipality-years were predicted with moderate (0.6-0.8) or strong (0.8-1.0) mean posterior predicted-class probabilities (i.e. P(Epidemic) for predicted epidemics and 1-P(Epidemic) for predicted non-epidemics), with over 70% of these moderate or strong predictions correct (Fig. S6). However, we also found that many misclassifiers “failed silently,” that is, outputted incorrect answers with high confidence^40^. Our results show that over half of missed true epidemics were predicted to be epidemics with less than 0.3 MPP, and likewise, over half of missed non-epidemics were classified as epidemics with over 0.7 MPP. In general, this implies that incorrect predictions were typically the result of strong model conviction against true outbreak status; that is, based on historical climate patterns, these municipality-years had periods of weather conditions conducive to either outbreaks or low dengue activity, but experienced the opposite. In a few cities that showed no strong evidence of weather signatures (i.e. Barueri, Rio de Janeiro; Fig. 4), mean posterior probabilities were more borderline (0.4-0.6), suggesting that the climatic distinction between epidemic and non-epidemic years may have been low in those locations, resulting in low separability in the data and higher occurrence of incorrect predictions.

We therefore endeavor to apply a loose typology to instances of misclassification, in order to better understand the limitations of the present modeling framework. For municipality-years whose epidemic status was misclassified with strong conviction (mean posterior predicted-class probability ≥ 0.8), it is likely that the dengue activity that year was highly anomalous given what had been experienced in historically-similar weather conditions (Fig. S7). For municipality-years whose epidemic status was misclassified with borderline conviction (i.e. mean posterior predicted-class probability close to 0.5), the error is more likely to be a consequence of insufficient data to discriminate between epidemic and non-epidemic years on the basis of weather patterns alone; that is, the models were not well suited to make this distinction in the first place (Fig. S7).

**Prediction results using SVD feature extraction approach**

*Correspondence with weather signatures*

Retrospective, out-of-sample forecasts trained on a yearly expanding window were produced for 10 years (2008-2017). Every year, the top predictive time intervals were automatically selected to participate in an ensemble voting system based on historical out-of-sample prediction accuracy and a accuracy threshold.

In this alternative system, the phenomenon of “weather signatures”, temporal regions of high predictive power within a municipality in Brazil, is also present. The SVD approach, although as not as frequent as the windowing method, successfully extracted relevant features from data for classification, as it can be seen in the weather signatures generated at Fig S6. Both approaches showed clear weather signatures for São Gonçalo, Manaus, Aracajú, and Juazeiro do Norte with less recognizable patterns for Guarujá, Três Lagoas, Maranguape and Barueri, while cities with poor performance exhibited no specific tendencies (Figs. 4, S6).

*Weather-based forecasting performance*

The overall accuracy of this approach was approximately 60% of all epidemic years across 20 municipalities in Brazil between 2012-2017, in contrast to the 72% accuracy of the method presented in the main text.

*Model performance by year*

During the last three years of the time series (2015-2017), epidemics were predicted by the weather-only models with 59% accuracy, with 12 of the 13 outbreaks in 2016 correctly forecasted. Conversely, non-epidemic years during 2014 and 2017 were particularly difficult to predict for non-epidemics, with only one and two of cities correctly forecasting non-epidemics for these years, respectively. Overall, 2016 was the most successful year, with 14 out of 20 correct predictions, followed by 2012 and 2015, with 13 out of 20 correct predictions, while 2014 and 2017 were the years with the worst performance, having less than 10 municipalities correctly predicted.

**Table S1. Population and land characteristics of 20 dengue-endemic study cities.**

| **City** | **Area (mi²)** | **Population*** | **Population Density (pp/mi²)** |
| --- | --- | --- | --- |
| Rio de Janeiro | 485 | 6320000 | 13030.93 |
| Belo Horizonte | 127.8 | 1433000 | 11212.83 |
| Aracajú | 70.22 | 571149 | 8133.71 |
| São Luís | 319 | 958,545 | 3004.84 |
| Sertãozinho | 155.6 | 101784 | 654.14 |
| Manaus | 4402 | 1793000 | 407.31 |
| Rondonópolis | 1608 | 144049 | 89.58 |
| São Gonçalo | 91.6 | 337273 | 3682.02 |
| Barra Mansa | 211.3 | 171125 | 809.87 |
| Eunápolis | 462 | 93413 | 202.19 |
| Tres Lagoas | 3941 | 96341 | 24.45 |
| Barueri | 24.78 | 240749 | 9715.46 |
| SaoVicente | 87.65 | 332445 | 3792.87 |
| Juazeiro do Norte | 95.97 | 249939 | 2604.35 |
| Parnaiba | 168.2 | 145705 | 866.26 |
| SantaCruz | 142.7 | 87582 | 613.75 |
| Maranguape | 228.1 | 113561 | 497.86 |
| Barretos | 604 | 112101 | 185.60 |
| Ji-Paraná | 2663 | 116610 | 43.79 |
| Guaruja | 55.44 | 290752 | 5244.44 |
| *city proper. Source: Demographic Statistics Database, United Nations Statistics Division 2010 | | | |

**Figure S1. Number of dengue fever epidemic years in Brazil, 2001-2015.** Data on annual cases for all municipalities in Brazil were available through 2015, shown here, and we obtained data separately through 2017 for the 20 study municipalities (black crossed circles).
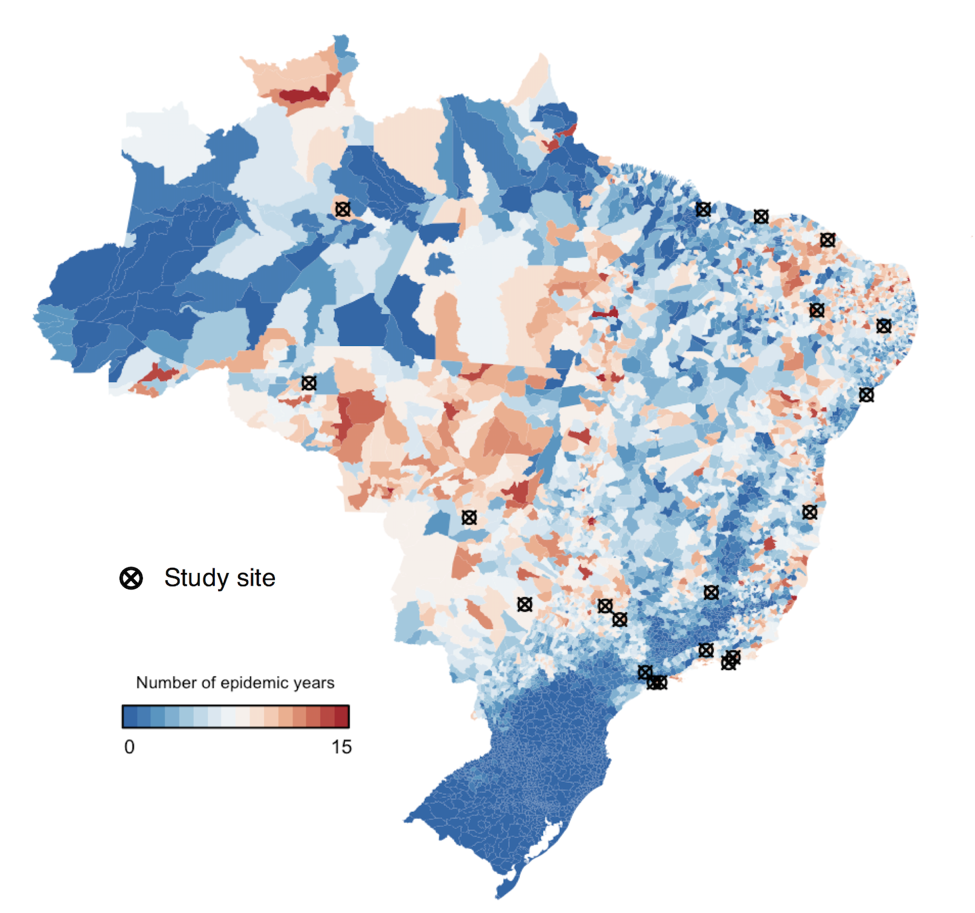

**Figure S2. Ensemble strength for 120 municipality-years (2012-2017)**. Ensemble strength is calculated for each of the 11 time windows selected into the ensemble each year, as a function of the historic out-of-sample accuracy of (a) the selected time window and (b) neighboring time windows (see *Supporting Information Materials & Methods)*. Municipalities are ordered by decreasing ensemble prediction accuracy; that is, the proportion of years correctly forecasted by the ensemble method over the years 2012-2017. Points are colored by prediction result (yellow=correct; green=incorrect).

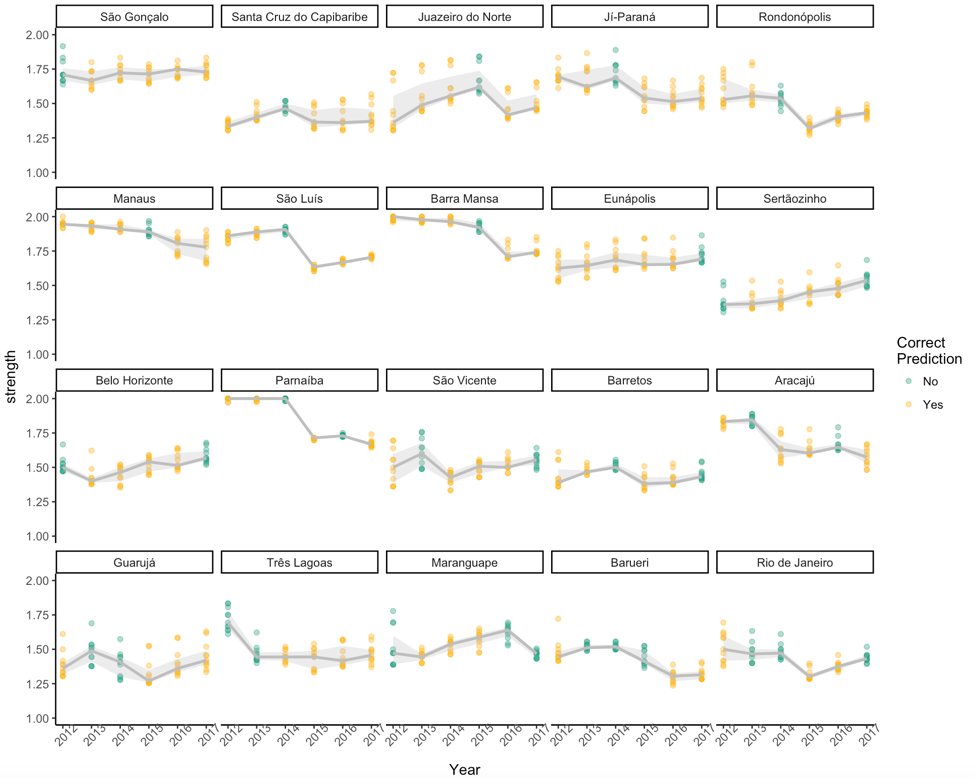

**Figure S3. Calibration curve of mean posterior probabilities over 120 municipality-years (2012-2017).**

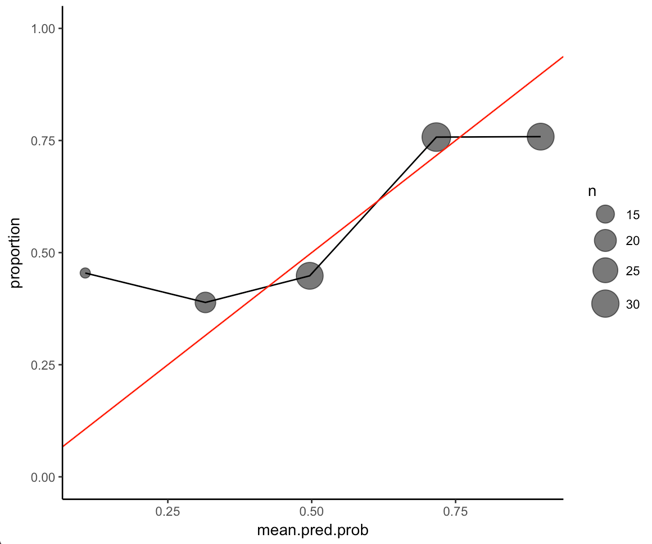

**Figure S4. The 10-year (2008-2017) out-of-sample forecast accuracy (%) for each time window of temperature and precipitation, by municipality, using Singular Value Decomposition feature extraction.** As described in *Supplemental Materials & Methods*, singular value decomposition was used as an alternative feature extraction method for comparison to taking the simple mean of weather-based variables over designated time windows, as presented in the main text. The x-axis (t_0_) indicates the start date of the time interval, and the y-axis (p) indicates the length of the time interval from which weather data were gathered (here, 10-99 days). achieving at least 7/10 correct out-of- sample forecasts are shown in shades of yellow. Municipalities are ordered by decreasing ensemble prediction accuracy; that is, the proportion of years correctly forecasted by the ensemble method over the years 2012-2017.

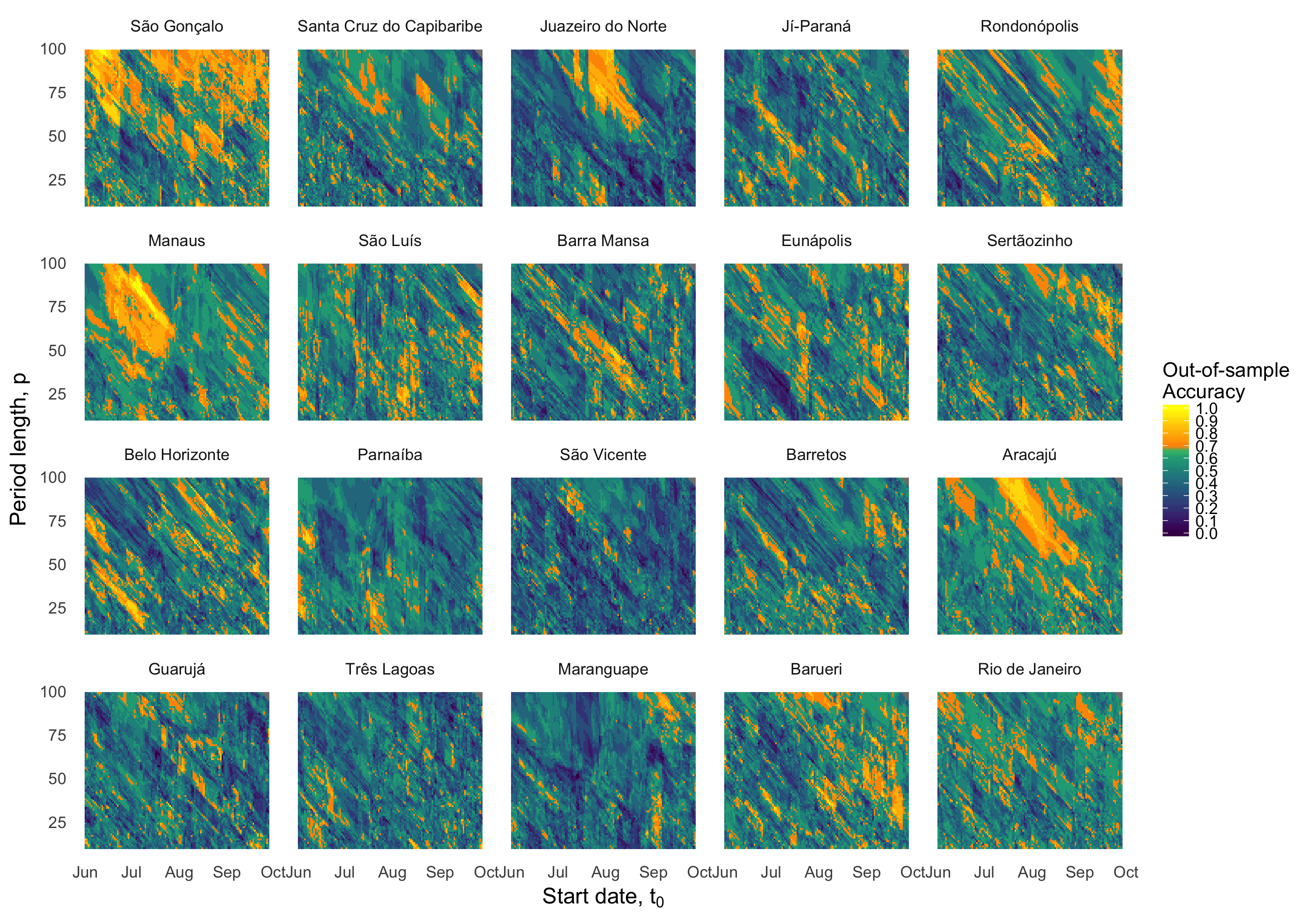

**Figure S5. Daily time series of weather inputs: 2000-2016 patterns of A) average temperature (K) and B) total precipitation (mm), by municipality.**

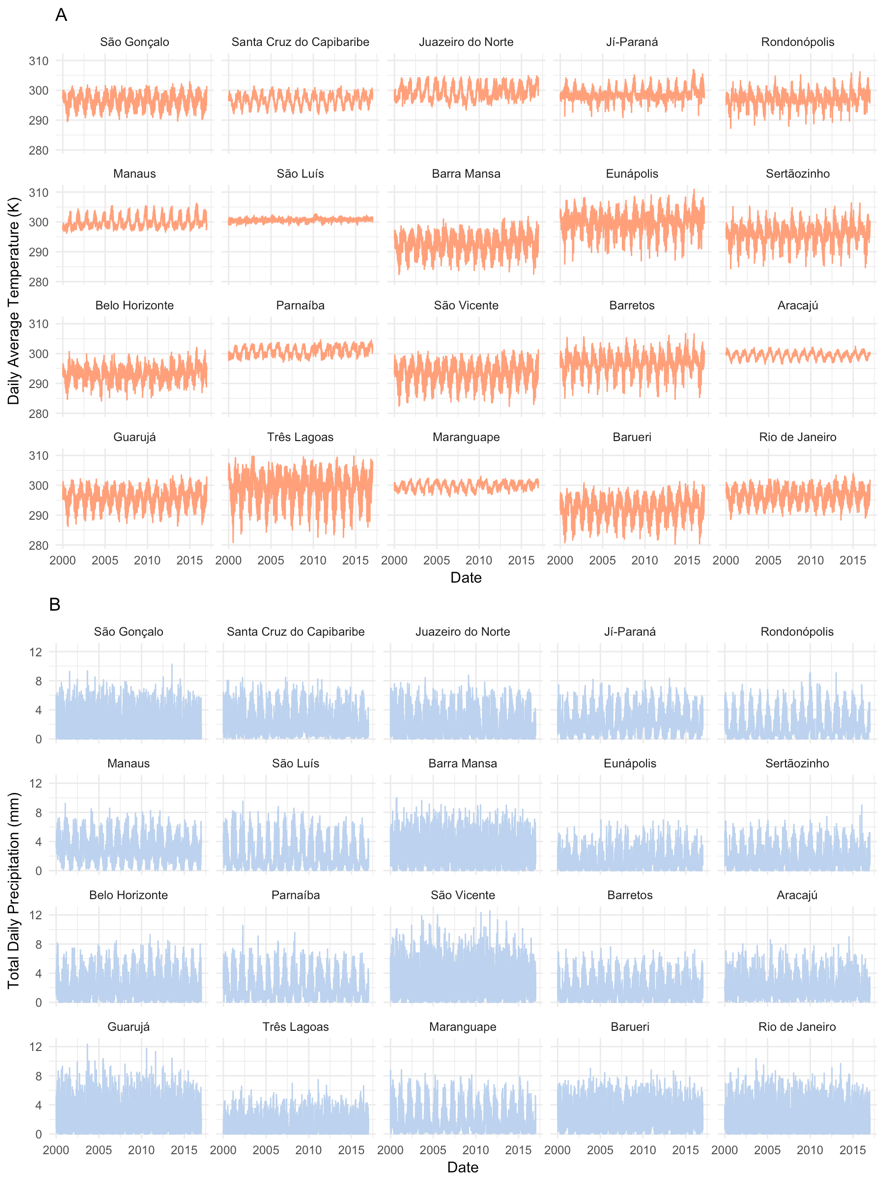

**Figure S6. Classifier and ensemble strengths.** Categories of classifier probability strength (weak: 0.2-0.4, borderline: 0.4-0.6, moderate: 0.6-0.8, and strong: 0.8-1.0) and ensemble strength (weak: 1.2-1.4, borderline: 1.4-1.6, moderate: 1.6-1.8, and strong: 1.8-2.0). The classifier probability is the mean posterior class probability, computed as P(Epidemic) for predicted epidemics and 1-P(Epidemic) for predicted non-epidemics, averaged over the 11 models of the ensemble. See *Supporting Information Materials & Methods* for calculation of the ensemble strength metric. There were no instances of probabilities < 0.2 nor of ensemble strengths < 1.2.

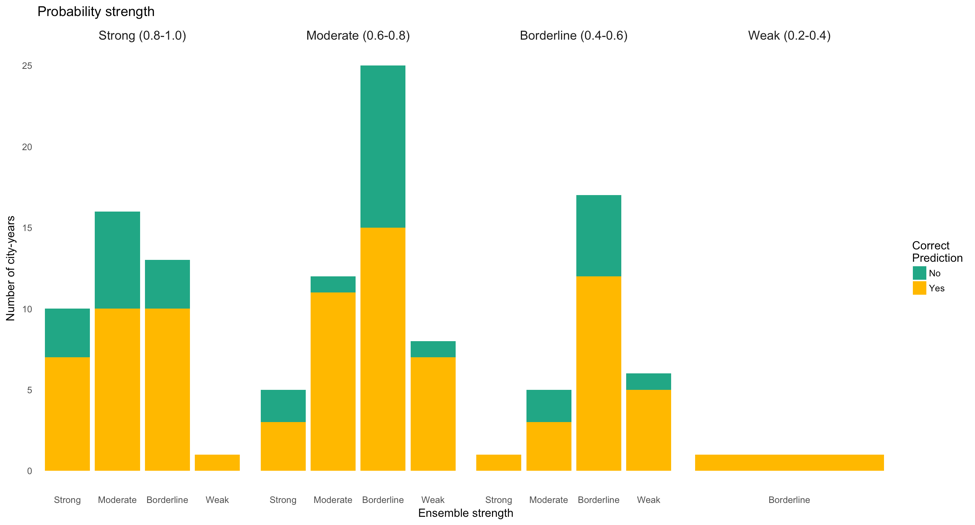

**Figure S7. Potentially anomalous or weakly-separable municipality-years for prediction.** (A) Predicted probabilities by municipality and year for ensemble forecasts (2012-2017). Predictions are colored by their true epidemic status (red=epidemic, blue=non-epidemic) with point shape indicating accuracy (closed circle=correct, cross=incorrect). A cyan circle designates potentially anomalous years, defined as years that were incorrectly predicted with strong conviction (mean posterior predicted class probability ≥ 0.8). A bright green circle designates years potentially following periods with low separability, defined as years that were misclassified with borderline conviction (0.4 ≤ mean posterior predicted class probability < 0.6).

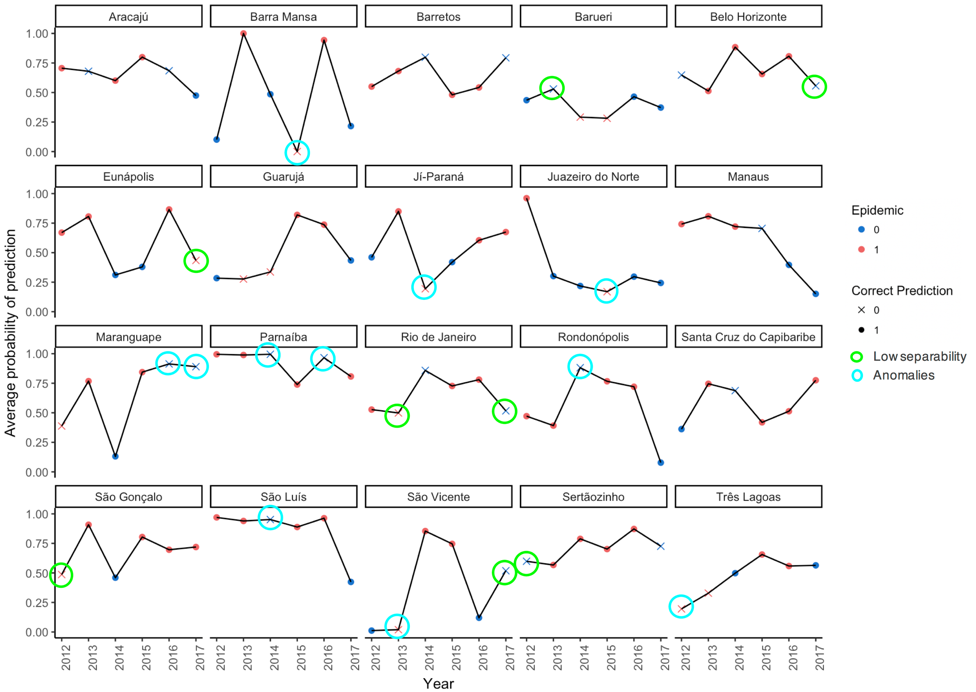
